# Machine Learning-based Prediction of the Long-term Stability of Chinese Hamster Ovary Cells due to Epigenetic Changes

**DOI:** 10.1101/2025.03.14.643414

**Authors:** Pedro Seber, Richard D. Braatz

**Author notes:** Contributing authors.

## Abstract

**Background:** Chinese hamster ovary (CHO) cells are the main system for producing recombinant protein biopharmaceuticals, but are inherently unstable, affecting their long-term productivity. This cell instability reduces their productivity over time during perfusion operation, which increases the costs of the resulting biopharmaceutical. No models have been published for the prediction of long-term stability.

**Results:** In this work, we create the first models for predicting the long-term stability of CHO cells due to changes in chromatin modification levels and methylation. Multilayer perceptrons are the best-performing models, reaching an F_1_ score of 59.1% and a Matthews correlation coefficient of 19.4%. The models are successful at identifying stable and highly productive CHO cells. Furthermore, Shapley values and interpretable models are used to investigate model coefficients, contributing biological insight to this problem and helping focus future data collection efforts. The models trained in this work are free and open source and available at github.com/PedroSeber/CHO_stability_prediction, allowing their use in the biopharmaceutical industry, reproduction of this work, and the retraining of models on other datasets.

**Conclusions:** We show that it is possible to train accurate machine learning models to predict the long-term stability of CHO cells using only epigenetic data. The models have high performance and excel in industrially relevant contexts, and thus can improve the bioproduction of medications, especially recombinant proteins. By providing the first predictive models for this task, this work also serves as a foundation for future data collection and modeling efforts.

## 1 Introduction

Biopharmaceuticals are an important class of drugs and were 66% of all new drug approvals by the U.S. FDA in 2017 [1, 2]. Chinese hamster ovary (CHO) cells are the main system for the production of recombinant protein biopharmaceuticals. In 2007, 70% of all recombinant protein biopharmaceuticals were produced in CHO cells [3], and this number increased to 84% in 2018 [1]. The predominance of CHO cells can be explained by their having many advantages for bioproduction, including high productivity, reaching titers > 10 g/l in the most productive scenarios [4]; amenability to being engineered, highlighted by how the average productivity of CHO cells increased 100-fold between the mid-1980s and mid-2000s [5]; that various human pathogens do not grow in CHO cultures [6]; that CHO cells can be cultured in serum-free suspension cultures [4], increasing their productivity, scale-up potential, and product consistency; and the ability of CHO cells to produce proteins that are properly folded and contain correct post-translational modifications [2].

To mass produce biopharmaceuticals, CHO cells are placed in bioreactors, typically in batch, fed-batch, or perfusion operation [7]. In batch operation, the culture receives no additions or removals until the end of the production run. Batch processes typically have the highest operating costs per g product, and typically have high batch-to-batch variation [8–10]. Fed-batch bioreactors are similar, but with nutrients added to the culture during the production run. Fed-batch bioreactors are the most common type used in industry, due to having higher productivity than batch while retaining relatively simple operation [11]. Both batch and fed-batch bioreactors require more labor to operate, as the cells need to be grown in bioreactors different from those used in the production phase.

After startup procedures are completed, perfusion bioreactors operate continuously, with media addition and bioreactor material removed at the same rate. The removed cells are retained in the bioreactor using a specialized membrane [7]. Perfusion bioreactors can have production runs that span multiple months [5], spending most of the time in quasi-steady-state operation, and lead to more consistent product quality [12]. Optimized perfusion bioreactors have higher volumetric productivity than fed-batch bioreactors, further reducing costs [7]. Also, biopharmaceuticals are delicate proteins, so quality-decreasing modifications, such as oxidation and aggregation, may occur while biopharmaceuticals are locked within a fed-batch bioreactor and away from ideal storage conditions [12, 13]. Perfusion reduces the effects of such phenomena by continuously removing the product from the bioreactor.

Despite these significant advantages, perfusion bioreactors have yet to dominate the industry. A critical challenge is CHO cell instability, which forces perfusion runs to be cut short, lowering their productivity and increasing costs. CHO cells are defined in the literature as being stable if they meet two criteria: retaining at least 70% of their volumetric productivity titer over at least 70 generations (population doublings) and generating a product with “no clinically meaningful differences” from the expected product [14]. Multiple causes of instability have been discovered and investigated, including genetic [15, 16], epigenetic, and culture-related issues. Epigenetic changes are a particularly important cause of instability [17–19]. Methylation, chromatin/histone modifications, and long non-coding RNA are the most impactful epigenetic modifications affecting stability [20–22]. In mammalian cells, the histone modifications H3K4me1 and H3K4me3 are commonly associated with transcriptionally active genes [23–25]; the modification H3K27ac is also associated with active enhancement of transcription [26]. Conversely, the modifications H3K27me3 and H3K9me3 inhibit transcription [27–30]. Finally, H3K36me3 is found within genes and is related to alternative splicing [31]. Further corroborating the importance of epigenetics, studies have shown that random or targeted (via CRISPR/dCas9) changes in methylation can lead to CHO clones with increased productivity or modified glycosylation [32–34].

In industry, the creation and selection of stable and productive CHO cells are performed completely empirically. The processes use methotrexate or methionine sulfoximine [5, 14, 35]. These chemicals are successful in leveraging natural selection to generate (relatively) productive CHO cells; however, it has been argued that the same chemicals also increase long-term genetic and epigenetic instability [5, 14, 35]. This selection procedure is also slow and expensive [5, 35], further increasing the challenges associated with obtaining stable and productive CHO cell lines. A significant limitation for the study and design of better CHO cell lines is that, although there are some models to predict immediate or short-term CHO cell productivities, there are no models for predicting long-term CHO cell stability. Ref. [36] sought to create a linear model for the productivity of CHO after 60 population doublings based on the levels of a few genes. However, that work used very few samples, does not report predictions on an independent test set (all metrics are reported for the train set), and has data quality issues; thus, the model is not usable in practice.

In this work, we construct machine learning models that use epigenetic data (histone modification and methylation levels in different locations of the genome) to predict the stability of CHO cells for biopharmaceutical production. These machine learning models are trained using efficient and open-source automatic machine learning (AutoML) software [37]. The best model, a multilayer perceptron (MLP) trained with the weighted focal differentiable MCC loss function [38], achieves an F_1_ score of 59.1% and a Matthews correlation coefficient (MCC) of 19.4% when predicting stability. When a “highly productive” class is extracted from the “stable” class to create a 3-class task, the best model achieves an F_1_ score of 42.4% and an MCC of 16.7%. Ablation tests show that histone modification data are sufficient to attain these metrics and that methylation data are not contributing to the predictive performance of the models.

## 2 Materials and Methods

This section describes the datasets and the model training methods. Instructions on running the AutoML software are provided on its GitHub repo (github.com/PedroSeber/SmartProcessAnalytics) [37].

### 2.1 Datasets

The stability data used in this work come from Ref. [39], which performed site-specific integration in many locations of the CHO genome and recorded the relative productivities of these CHO cells after 36 and 72 population doublings. This dataset is the largest, most recent compendium of long-term CHO cell stability data. These data also include the specific location where the integration was performed and the DNA nucleotide bases of that location. 4,783 samples generated via Piggybac cloning are used for 5-fold cross-validation (CV), and 550 samples generated via lentivirus integration are used as a test set. Different integration methods were separated into these two sets to ensure that the model can predict long-term stability in a way that is agnostic to the integration method used. This work does not consider whether the similarities and differences between integration preferences of different methods [40, 41] play a material role in predicting long-term stability of CHO cells. Lentivirus integration was used as the test set due to its smaller number of samples. The number of samples in each set is slightly smaller than the total samples found in Ref. [39] because samples had to be cleaned up for use with epigenetic data.

Originally, this work sought to model this numerical relative productivity as a function of the DNA bases of the integration site; however, that approach failed to yield any useful models (test *R*^2^ *→* 0). The use of epigenetic data (described below) with the same targets led to similar results, and this issue is discussed further in Section A2 of the Appendix, which also features an analysis of the productivity data. To remedy this problem, the numerical productivity data were stratified into “stable” and “unstable” classes. As per the FDA definition (Section 1, [14]), a sample was labeled as “stable” if its relative productivities after 36 and 72 population doublings were *≥* 0.7; else, it was labeled as “unstable.” For the 3-class problem, a sample was labeled as “highly productive” if its relative productivities after 36 and 72 population doublings were *≥* 2, which is equivalent to selecting the 5% most productive samples. The raw epigenetic data used in this work come from and are described in Ref. [19]. These data were preprocessed using bespoke code written that leveraged commonly used tools including BWA-MEM [42], samtools [43], and MACS3 [44]. The main preprocessing steps include realignment to the CriGri-PICRH-1.0 genome and peak calling, and are detailed in the code files within the GitHub repository. The processed data are comprised of the levels of different histone modifications at different time points (Tp, as defined by Ref. [19]) and the methylation levels of the bases surrounding an integration site. There are 97 chromatin/histone features and a varying number of methylation features (depending on the window size, a hyperparameter).

The (lack of) effect of the methylation window size are discussed in Appendix A3.

### 2.2 Data-driven model creation and evaluation

Logistic regression (LR), logistic regression with the elastic net penalty (EN) [45], random forest (RF) [46], support vector machine with radial basis functions (SVM) [47], LASSO-Clip-EN (LCEN) [48], and MLP models trained with the cross-entropy (MLP-CE) and weighted focal differentiable MCC [38] (MLP-diffMCC) loss functions are trained on the data described above using the AutoML software. Within that software, LR, EN, RF, and SVM models are constructed with scikit-learn [49], LCEN models are constructed as per [48], and MLP models are constructed with PyTorch [50]. A list of the hyperparameters used for each model type is available in Section A1.

The best combination of hyperparameters for each model and task is determined by grid search, and the combination with the highest cross-validation average F_1_ score is selected. F_1_ scores and MCCs on an independent test dataset are then obtained. The F_1_ score is used due to its efficacy in balancing recall and precision (two important metrics), its widespread usage, and its flexibility for prioritization using a different *β* value. The MCC is used because it is widely regarded as the single best metric for classification tasks [51, 52]. Further description and explanation of these metrics is available in Section 2.3 of Ref. [53]. This procedure is repeated three times so that a mean *±* standard deviation of test errors may be reported.

### 2.3 Interpretability studies

Interpretability studies are performed in Section 3.3 by obtaining scaled coefficients for the MLP-diffMCC and LCEN models. The magnitude of the scaled coefficients predicted by a model can be used as a proxy for the importance of a feature. For interpretable models (such as produced by LCEN), the coefficients may be obtained directly from the model. For non-interpretable models (such as produced by MLPs), a versatile yet powerful method to obtain these coefficients is using an *a posteriori* interpretability method, such as Shapley values [54]. In this work, Shapley values are calculated using the shap Python package [55].

## 3 Results

### 3.1 MLP models predict stability as a function of histone modification levels with good performance

This section includes the results of the models trained with the 2-class version of the stability data as a function of chromatin modification levels. Multiple model types (as described in Section 2.2) were trained for this task. The LR and EN models had very similar performance, slightly above baseline (Fig. 1 and Table 1), where the baseline is not using any model. These models were average relative to all models tested in this work. The SVM and RF models typically had lower performances despite their nonlinear classification abilities. RF achieved medium-high MCCs relative to other models at some thresholds (Fig. 1). LCEN achieved the highest performance out of all non-deep learning models, reaching an F_1_ score of 53.8% at its best threshold and an MCC of 7.54% at that threshold. Nevertheless, that performance is low overall. Both types of MLP models performed better, and those trained with the weighted focal differentiable MCC achieved the highest performance (Fig. 1 and Table 1). At their best threshold, the MLP-diffMCC models reached an F_1_ score of 59.1% and an MCC of 19.4%. These results indicate that nonlinear models are needed to predict the stability of CHO cells; deep learning models, which are universal function approximators, obtained the best predictive performance. We hypothesize that the main reasons that prevent the construction of models that have even higher predictive performance involve the nature of the data (chromatin is not the only biological mechanism of instability, but no dataset with other features is available) and the quality of the stability data (as elaborated in Section A2).

**Fig. 1:**
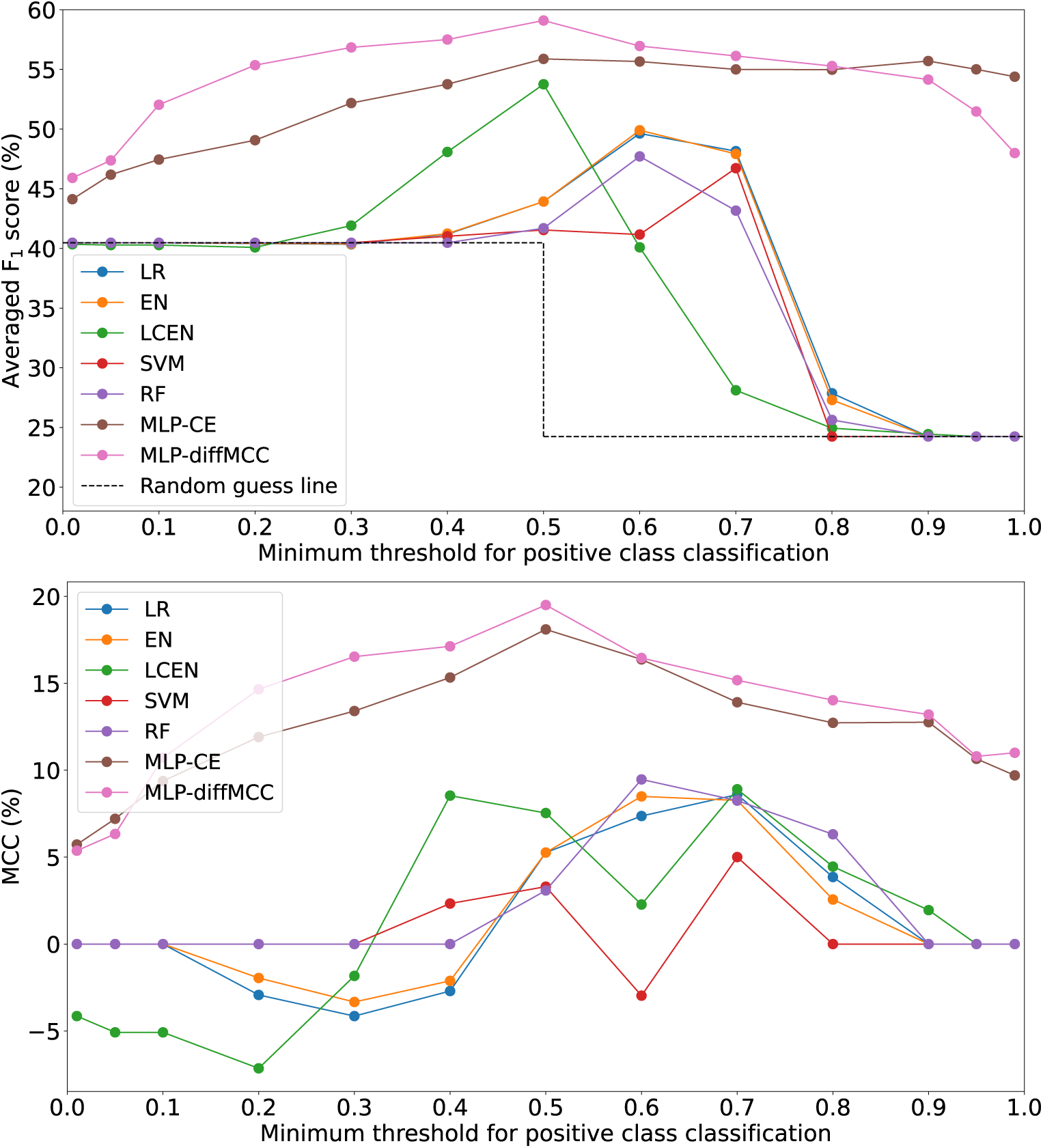
Averaged F_1_ scores (top) and Matthew correlation coefficients (MCCs) (bottom) for the models trained with the 2-class version of the stability data as a function of chromatin modification levels. Model labels are as in Section 2.2. “Random guess line” shows the F_1_ score obtained by guessing all samples as positive or negative.

**Table 1:** Test-set averaged F_1_ scores and MCC for different models trained with the 2-class version of the stability data as a function of chromatin modification levels. Model labels are as in Section 2.2. “No Model” refers to the F_1_ score obtained when all samples are guessed as positive and is the minimum value any model should achieve (a baseline). The highest metrics are highlighted in bold.

| Model | Averaged $F_1$ Score (%) | MCC (%) |
| --- | --- | --- |
| LR | 49.6 | 7.37 |
| EN | 49.9±0.3 | 8.50±1.22 |
| LCEN | 53.8±0.2 | 7.54±0.48 |
| SVM | 46.7±0.5 | 5.01±0.68 |
| RF | 47.7±1.7 | 9.47±0.20 |
| MLP-CE | 56.4±1.3 | 17.1±2.3 |
| MLP-diffMCC | **59.1±0.9** | **19.4±1.1** |
| No Model | 40.5 | 0 |

Although the preprocessed dataset also contained methylation data for the samples (Section 2.1), these data were not included in the input due to a lack of effect on the predictive performance of the models (Fig. 2; further elaborated in Section A3). Paired t-tests comparing the F_1_ scores and MCCs of the models trained without and with methylation data (for a window size = 20) returned p-values equal to 0.797 and 0.349, respectively.

**Fig. 2:**
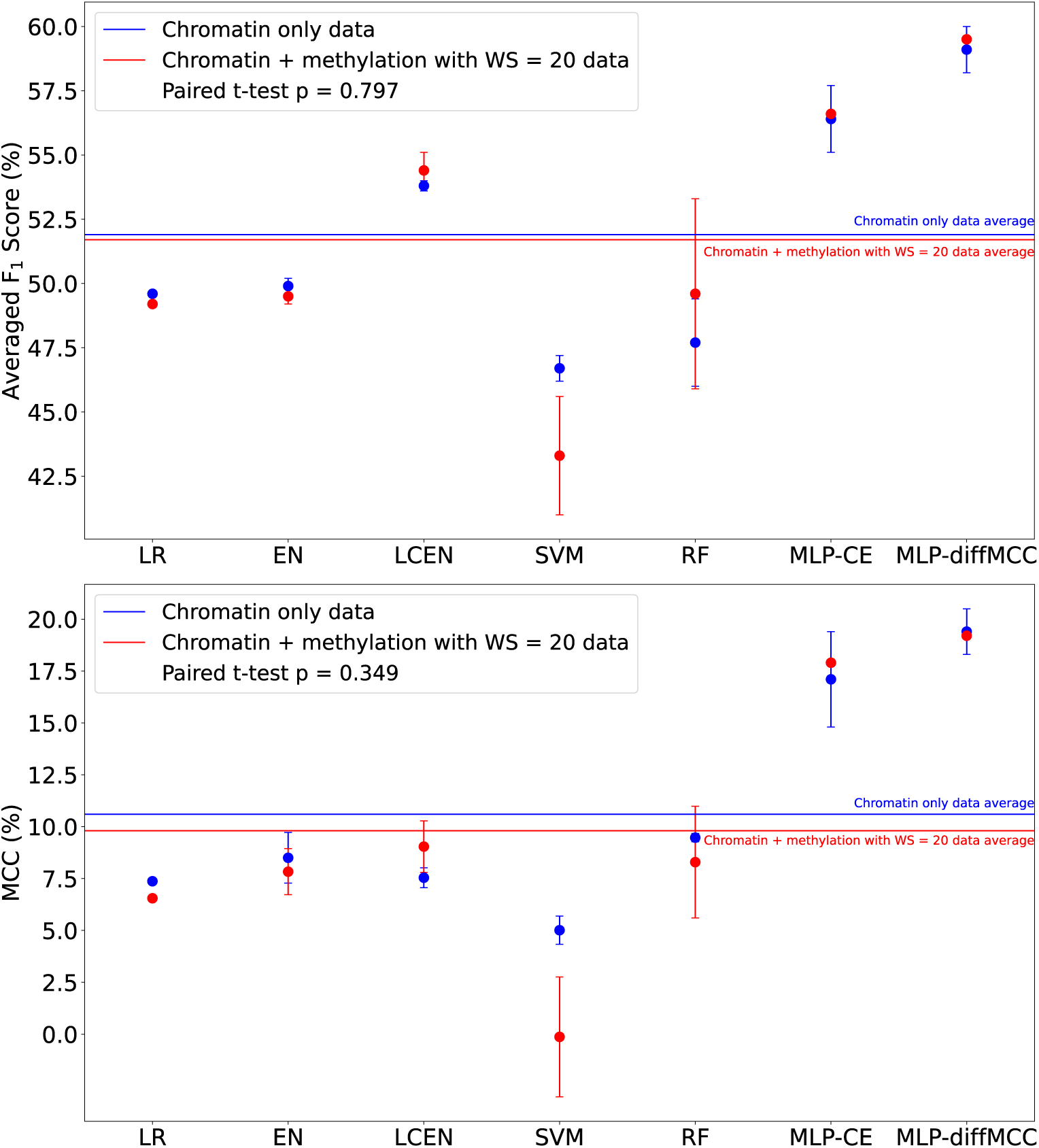
F_1_ scores (top) and Matthew correlation coefficients (MCCs) (bottom) for the models trained with the 2-class version of the stability data as a function of chromatin modification levels (blue) or both chromatin modification levels and methylation with a window size of 20 (red). Model labels are as in Section 2.2. Error bars are sample standard deviations (*n* = 3).

### 3.2 Prioritizing industrially relevant outcomes with the F*_β_* score

As mentioned in Section 2.2, models and their hyperparameters were selected based on their averaged F_1_ score. That same averaged F_1_ score metric is widely reported in the results of this work (Sections 3.1 and 3.4). This metric was chosen due to its universality, which leads to the creation of models that are widely applicable, especially to those interested in the biological mechanisms behind CHO cell instability.

However, the averaged F_1_ score may be of less interest to the biopharmaceutical industry. In industry, due to financial and temporal costs and restrictions, it is much less important to detect all stable CHO clones or to focus on unstable CHO clones. Instead, industry focuses on finding just enough stable clones but demands that each of these clones is accurately labeled as positive. Misidentification of an unstable clone as stable (a false positive) can lead to significant problems, including losses in revenue and regulatory issues, but misidentification of a stable clone as unstable (a false negative) is much less consequential. Thus, in an industrial context, other metrics are more relevant than the averaged F_1_ score.

The F_1_ score is part of a family of metrics called the F*_β_* score,

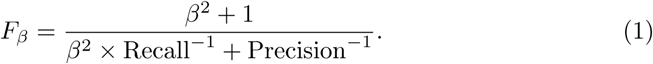

These scores are weighted harmonic means of the recall and precision, such that the *β* parameter indicates how many times recall is weighted over precision.^1^ The use of different *β* increases the flexibility of the F*_β_* score metric and allows for prioritization of goals (as mentioned in Section 2.2). In the industrial context described above, false positives are much more consequential than false negatives; thus, precision is more important than recall, and so F*_β_* scores with *β <* 1 are a more relevant metric than the F_1_ score. In addition, F*_β_* scores of the positive class are more relevant than the averaged F_1_ score, as the performance on predicting the unstable class is less relevant for the biopharmaceutical industry.

The MLP-diffMCC models from the previous section are evaluated in terms of their F*_β_* scores for 0 *≤ β ≤* 1. These models are selected because they had the highest performance among all models tested (Fig. 1 and Table 1). Because these models had higher precision than recall at most thresholds, the performance of MLP-diffMCC models consistently increases as *β* is lowered (Fig. 3). In addition, we evaluate the performance of these models at a threshold of 0.99, as high thresholds reduce false positives at the expense of increasing false negatives, using the positive-class F*_β_* score. The performance of the MLP-diffMCC models increases with decreasing *β* and becomes more consistent (lower standard deviations) (Fig. 4). When *β* = 0, the models reach a positive-class F_0_ score (precision) = 74.9% *±*1.8, indicating that three out of four positive predictions made by the model are correct. This high performance is essential for the adoption of these models by the biopharmaceutical industry, which can decrease drug costs and accelerate production timelines. Finally, it is important to keep in mind that these models were trained and had their hyperparameters selected to maximize their positive-class F_1_ scores, yet attain high performance in this condition with different target metrics. Models can be trained using the same methodology and data to maximize their precision or a certain F*_β_* score, which will lead to even higher metrics for this industrially relevant scenario.

**Fig. 3:**
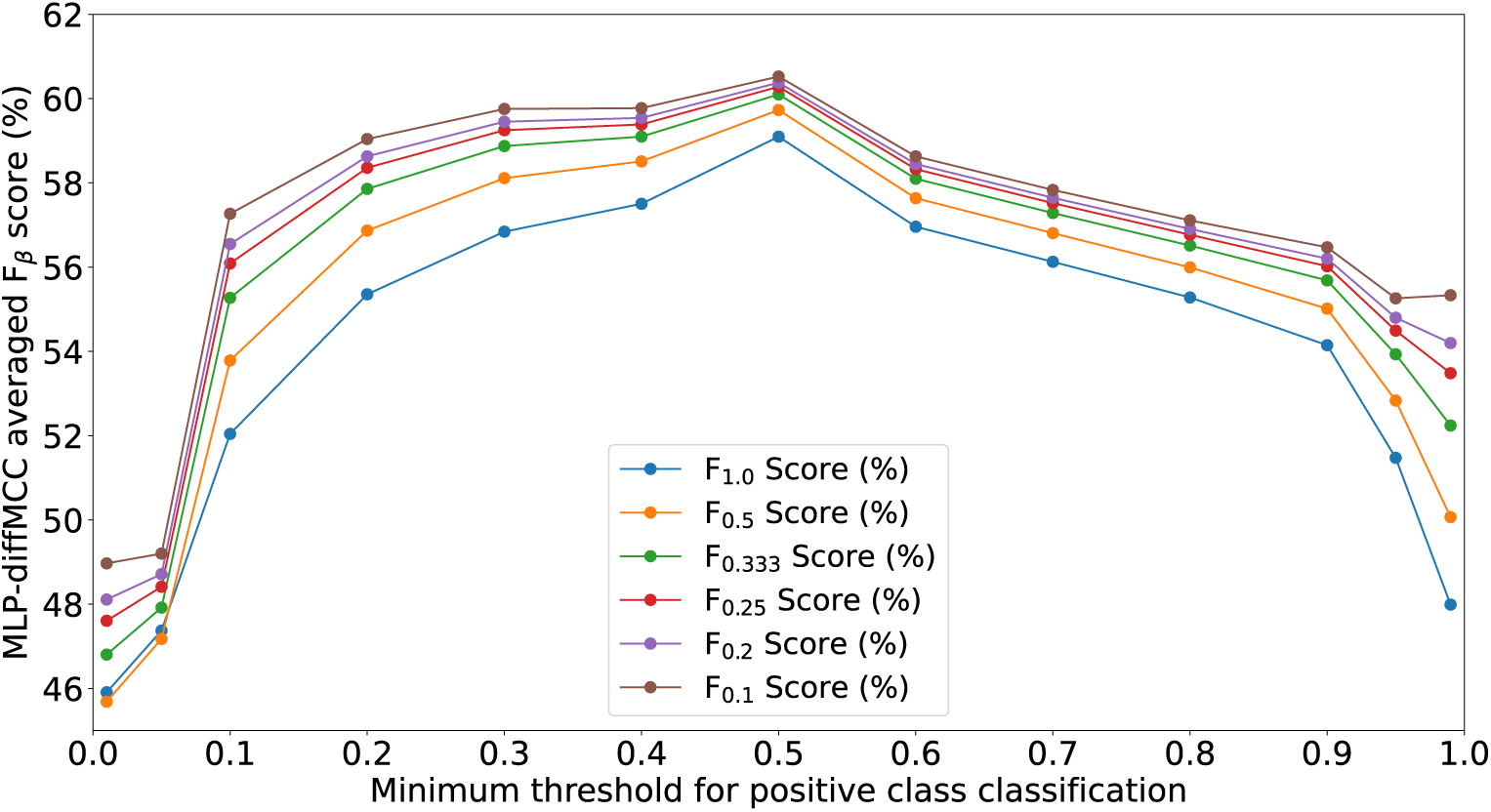
Averaged F*_β_* scores for the MLP-diffMCC models trained with the 2-class version of the stability data as a function of chromatin modification levels. The F_1_ score shown here is the same curve as in Fig. 1 (top) for the MLP-diffMCC models.

**Fig. 4:**
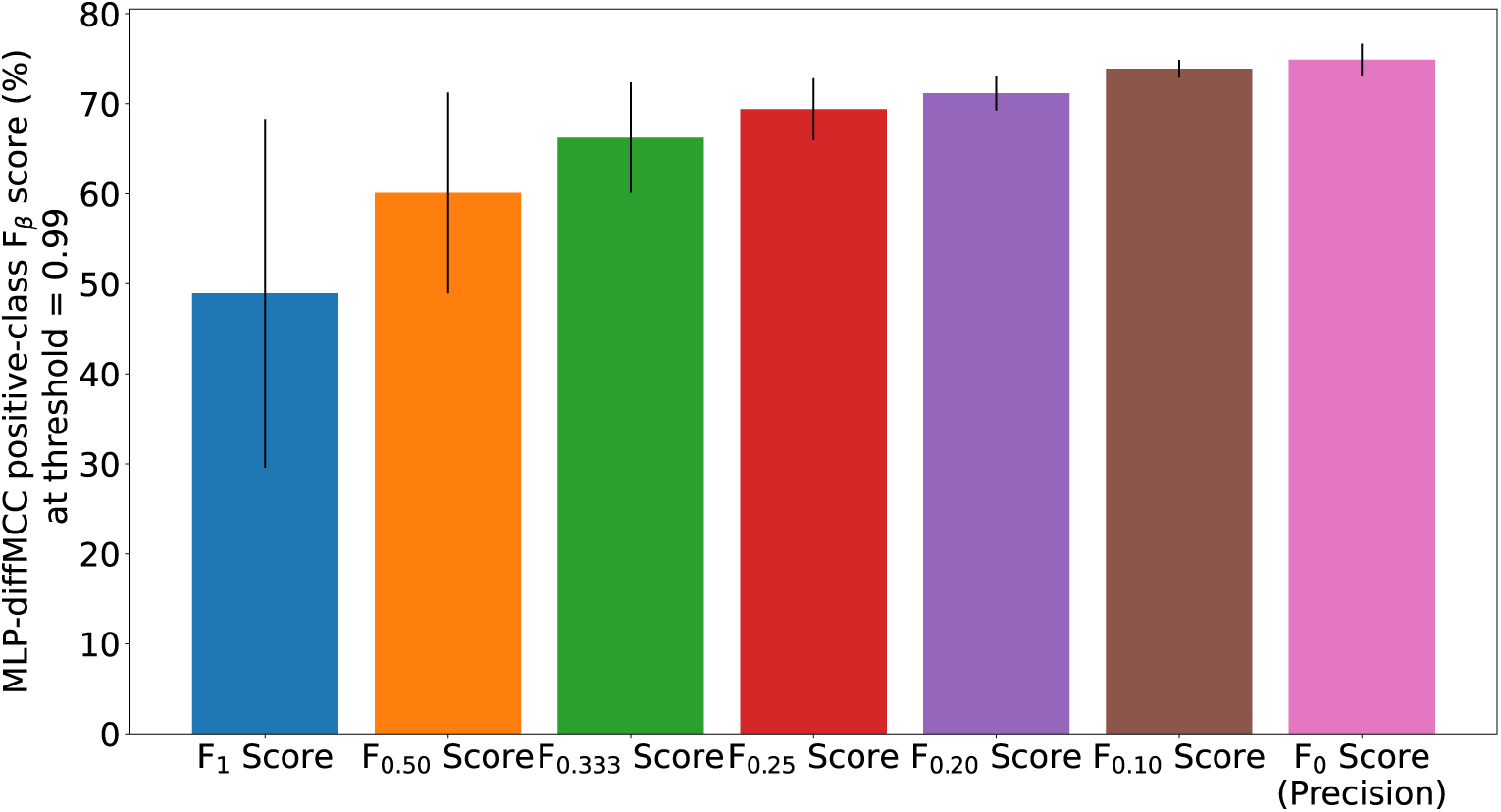
Positive-class F*_β_* scores for the MLP-diffMCC models trained with the 2-class version of the stability data as a function of chromatin modification levels. Black lines on each bar are sample standard deviations.

### 3.3 Interpreting the model coefficients to obtain biological insight

Coefficients for all seeds of the LCEN and MLP-diffMCC models were obtained, and the features corresponding to the 20 largest coefficients (in absolute magnitude) were analyzed to determine how often they are shared among different seeds of the same model (Fig. 5). Multiple types of sharing were considered. In decreasing order of exactness of match: the best-case scenario of 3 exact matches (purple in Fig. 5), 2 exact matches and a third off by one timepoint (Tp) (green), 2 exact matches (cyan), 3 matches but off by one Tp among the models (yellow), 2 matches but off by one Tp (red), and no matches (black). To serve as a baseline and to determine whether any sharing done by the models is meaningful, sets of random numbers were generated 10,000 times, and the same types of “feature sharing” were analyzed among these random sets. In this random baseline, about 55% of all top-20 features are not shared or only poorly shared (red and black sectors of Fig. 5, right). Conversely, MLP-diffMCC models have good sharing for over 58% of their features (purple, green, and cyan sectors of Fig. 5, left), and this number increases to two-thirds of the features for the LCEN models (same sectors, Fig. 5, center). This high rate of feature sharing indicates that the models are consistently learning to focus on the same components, highlighting the robustness and reproducibility of this approach. Chi-squared tests comparing the sharing of features from the models vs. that of random features return a p-value of 2.9*×*10*^−^*^7^ for the MLP-diffMCC models and 8.3*×*10*^−^*^10^ for the LCEN models, demonstrating that this high rate of feature sharing is very unlikely to be a coincidence.

**Fig. 5:**
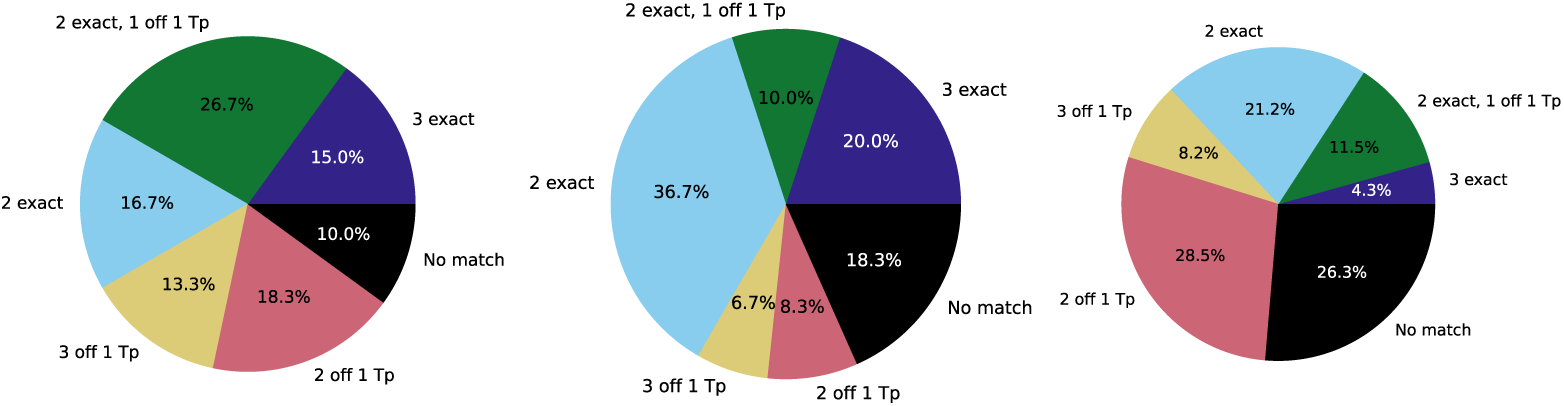
Proportion of features shared among the top-20 features of MLP-diffMCC (left) and LCEN (center) models, with sets of random features (right) used as a baseline. The types of matches considered, in decreasing order of exactness of match, are: the best-case scenario of 3 exact matches (purple in Fig. 5), 2 exact matches and a third off by one timepoint (Tp) (green), 2 exact matches (cyan), 3 matches but off by one Tp (yellow), 2 matches but off by one Tp (red), and no matches (black).

The histone modifications shared most often among all 6 models and seeds were H3K27ac, H3K4me1, and H3K4me3 (at specific Tp). As mentioned in Section 1, H3K4me1, H3K4me3, and H3K27ac are commonly associated with transcriptionally active genes and enhancement of transcription. As such, the models are not only learning consistent features, but also features that have biological relevance to this stability prediction problem. These results can assist in providing biological insights into the epigenetic mechanisms of CHO cell instability and assist future data collection efforts.

### 3.4 MLP-diffMCC succeeds on the challenging prediction of highly productive clones

As described in Section 2.1, samples with relative productivity *≥* 2 were split into a third class^2^ to determine whether the models could correctly identify these samples, as they are of the highest interest to the biopharmaceutical industry. Once again, only chromatin data were used for these predictions, and the same models used in Section 3.1 were used here. This task was significantly more challenging than the 2-class problem, in part because of the low abundance of highly productive samples (only 5% of the training set samples were highly productive). Once again, the LC and EN models had very similar, slightly below-average performances (Table 2). SVM had the lowest performance overall, even resulting in negative MCCs for some seeds. RF was only marginally better than these other models, and could not attain the higher MCCs (when compared to other models) that it attained in the 2-class task (Section 3.1). Although LCEN had the best performance among the non-MLP models, it still had a lower performance than in the 2-class task. The MLP models were again the best-performing models. The MLP-CE models had a considerably higher MCC than the non-MLP methods, but their F_1_ scores were comparable to those from LCEN models. These results contrast those from the 2-class task, in which MLP-CE slightly outperformed LCEN even in F_1_ score terms. Finally, the MLP-diffMCC models displayed the highest metrics on this task, reaching an F_1_ score of 42.4% and an MCC of 16.7%. These metrics are slightly lower but comparable to those from the 2-class task, especially in terms of MCC.

**Table 2:** Test-set averaged F_1_ scores, MCC, and “highly productive” class F_1_ scores for different models trained with the 3-class version of the stability data as a function of chromatin modification levels. Model labels are as in Section 2.2. “No Model” refers to the F_1_ score obtained when all samples are guessed as positive and is the minimum value any model should achieve (a baseline). The highest metrics are highlighted in bold.

| Model | Averaged $F_1$ Score (%) | MCC (%) | Highly productive class $F_1$ score |
| --- | --- | --- | --- |
| LR | 29.1 | 2.57 | 0 |
| EN | 29.1±0.0 | 2.57±0.00 | 0±0 |
| LCEN | 35.5±0.4 | 5.53±0.20 | 9.95±0.65 |
| SVM | 28.1±1.9 | 0.11±2.07 | 0±0 |
| RF | 29.9±2.3 | 3.12±1.28 | 0±0 |
| MLP-CE | 37.1±2.1 | 13.4±0.5 | 9.03±2.27 |
| MLP-diffMCC | **42.4±0.2** | **16.7±0.7** | **15.1±2.1** |
| No Model | 26.4 | 0 | 0 |

A significant issue with the non-LCEN, non-MLP models is that they all had a test-set highly productive class F_1_ score of 0 (Table 2), indicating that they did not correctly classify any of the test samples as highly positive. This lowers these models’ average metrics and makes them effectively useless for this 3-class predictive task. The MLP-diffMCC models had a highly productive class F_1_ score equal to 15.1%, the highest of the models. These results further corroborate the idea that the stability of CHO cells is a highly complex and nonlinear function of chromatin modification levels and that this complexity is increased when a third, infrequent class is added to the dataset.

## 4 Discussion

This work constructs machine learning models to predict the long-term stability of CHO cells based on epigenetic data: chromatin modification and methylation levels. Multiple methods are cross-validated and tested for this task (Section 2.2). The data used in this study were published in different works [19, 39] and preprocessed and merged by us. Preprocessing steps included converting the stability values to classes and transforming the raw ChIP-Seq data into usable values (Section 2.1). Although the merger of two different datasets is not ideal, since the input data (epigenetic features) come from cells different from those that originated the output data (relative productivity over time, which was used to determine long-term stability), this merger is necessary because no public dataset collected both (or similar) data. Ref. [39] did not collect epigenetic or bioreactor data, and Ref. [19] did not collect long-term CHO cell stability data. This lack of publicly available, high-quality data limits the performance of data-driven models to predict long-term CHO cell stability. Nevertheless, in this work, we show that such predictive models can be constructed even with the data limitations that exist.

The first task explored in this work was a 2-class prediction of whether a CHO cell is stable in the long term based on its chromatin modification levels (Section 3.1). All of the models were better than the baseline, and the best-performing models were (respectively) MLP-diffMCC, MLP-CE, and LCEN (Table 1 and Fig. 1). These models are all nonlinear and can model complex functions, corroborating the idea that the stability of CHO cells is determined by intricate processes. As shown in Fig. 1, the MLP-diffMCC achieved the best metrics at almost every threshold. At its best threshold, the MLP-diffMCC model reached an averaged F_1_ score of 59.1% and an MCC of 19.4%. We hypothesize that data quality issues could be limiting the performance of the models. A surprising result was that adding methylation data to this dataset did not improve the models’ predictive performances (Fig. 2 and Section A3), which we hypothesize occurs because there are very few methylated residues overall in the dataset (Table A2). We show that these models are also successful at predicting stability in contexts that are more relevant to the biopharmaceutical industry (Section 3.2). In this context, precision is more relevant than recall; thus, F*_β_* scores with *β <* 1 are more important, especially the positive-class F*_β_* score. The same MLP-diffMCC models can reach a precision of 74.9% (Fig. 4), highlighting their usability also in industrially relevant scenarios.

The 20 largest coefficients of all seeds of the MLP-diffMCC and LCEN models were also analyzed for interpretability and biological insights (Section 3.3). Models were sharing a majority of their features, and at a much higher frequency than expected by chance (Fig. 5). The features shared among all models and seeds were histone modifications associated with transcription activation (H3K4me1, H3K4me3, and H3K27ac) and can be an important direction for future studies or data collection. These results show that the models are not only consistently learning using the same features, but also that the features being learned are biologically relevant, increasing the robustness of and trust in the models from this work.

Next, a “highly productive” class was extracted from the “stable class” to form a 3-class dataset (as described in Section 2.1). This “highly productive” class is of great interest to the biopharmaceutical industry, so models that can successfully classify it have high commercial and practical value. Most models struggled on this task, in part because of the low abundance (5%) of highly productive samples (Section 3.4 and Table 2). Critically, most models had a test-set “highly productive” class F_1_ score of 0, indicating they did not classify any samples of that class correctly. The LCEN and MLP models succeeded in this task, although to a lesser extent than in the 2-class task. The biggest exception were the MLP-diffMCC models, which maintained most of their MCCs (16.7%) and F_1_ scores (42.4%) in this task when compared to their 2-class metrics. Furthermore, MLP-diffMCC models had the highest test-set “highly productive” class F_1_ score, equal to 15.1% (Table 2).

Overall, these modeling results show, for the first time, the ability to predict the long-term stability of CHO cells. As mentioned in Section 1, successfully predicting this stability is essential for using CHO cells in perfusion bioreactors, which are more productive, and can lower the costs of biopharmaceuticals. We expect that the models in this work can be used by the biopharmaceutical industry to guide the discovery of stable CHO cell clones. The use of modeling can also considerably reduce the time needed for the identification of stable CHO to just under 10 days. Furthermore, the interpretability studies (Section 3.3) can assist future studies or data collection efforts. The use of AutoML methods makes the approach accessible even to non-experts in machine learning and facilitates the use of these models in industrial contexts. Finally, we freely release the models trained in this work, allowing their direct use for research or bioproduction. Future work could collect more accurate data and integrate other features, such as genetic or bioreactor culture information, into this dataset.

## Availability of data and materials

The processed data, code used to process these data, and MLP-diffMCC models from this article are available in a GitHub repository at https://github.com/PedroSeber/CHO_stability_prediction.

## Competing interests

The authors declare that they have no competing interests.

## Funding

Financial support is acknowledged from the U.S. Food and Drug Administration (75F40121C00090). PS was supported by a Takeda Fellowship.

## Author contributions statement

P.S.: conceptualization, methodology, data curation, software, validation, formal analysis, writing – original draft, writing – review & editing, visualization.

R.D.B.: conceptualization, resources, writing – original draft, writing – review & editing, supervision, project administration, funding acquisition.

## Appendix A1 List of hyperparameters used in this work

All possible combinations of the hyperparameters below were cross-validated.

1. For the logistic regression with elastic net penalty (EN) models: *α* = 0 and 20 log-spaced values between *−*4.3 and 0 (as per np.logspace(-4.3,0,20)) and L_1_ ratios = [0, 0.1, 0.2, 0.3, 0.4, 0.5, 0.6, 0.7, 0.8, 0.9, 0.95, 0.97, 0.99].
2. For the LCEN models: *α* and L_1_ ratios as above, *degree* values = [1, 2], *lag* = 0, and *cutoff* values between 8*×*10*^−^*^4^ and 1*×*10*^−^*^2^.
3. For the random forest (RF) models: [10, 25, 50, 100, 200, 300] trees, maximum tree depth = [2, 3, 5, 10, 15, 20, 40], minimum fraction of samples per leaf = [0.01, 0.02, 0.05, 0.1], and minimum fraction of samples per tree = [0.1, 0.25, 0.333, 0.5, 0.667, 0.75, 1.0].
4. For the AdaBoost (AdaB) models: [10, 25, 50, 100, 200, 300] trees/estimators and learning rates = [0.01, 0.05, 0.1, 0.2].
5. For the support vector machine (SVM) models: C values = [0.01, 0.1, 1, 10, 50, 100], epsilon values = [0.01, 0.025, 0.05, 0.075, 0.1, 0.15, 0.2, 0.3], and gamma values = [1/50, 1/10, 1/5, 1/2, 1, 2, 5, 10, 50] divided by the number of features in a dataset were used.
6. For the multilayer perceptron (MLP) models: representing an MLP with one hidden layer as [X] and an MLP with two as [X, Y], hidden layer sizes of {[97], [194], [291], [388], [485], [582], [97, 97], [194, 194], [194, 97], [291, 291], [291, 194], [388, 388], [388, 291], [485, 485], [485, 388], [582, 582], [582, 485]} were used.

Learning rates = [0.05, 0.1, 0.5] (and also 0.01 in the 3-class problem), the ReLU activation function, batch sizes of [512, 1024, 2048], 100 epochs, and a cosine scheduler with a minimum learning rate equal to 1/16 of the original learning rate with 10 epochs of warm-up. For the weighted focal differentiable MCC loss function, gamma values = [1.5, 2.0] were also used. No loss-function class weights were used in the 2-class problems. Loss-function class weights of {[1, 1, 1], [1, 1, 2], [2, 1, 2], [2, 1, 3], [2, 1, 4], [2, 1, 5], [2, 1, 6], [2, 1, 7], [2, 1, 8], [2, 1, 10], [3, 1, 5]} were used in the 3-class problems.

## Appendix A2 Comments on the numerical relative productivity data

As mentioned in Section 2.1, the numerical relative productivity data from Ref. [39] failed to yield any useful models (test *R*^2^ *→* 0) no matter whether genetic or epigenetic data were used. An analysis of the properties of these productivity data (Table A1) reveals some unexpected patterns, which include that (1) hundreds of samples are stable at *t* = 72 but not at *t* = 36, (2) about half the samples in each set are more productive at *t* = 72 than at *t* = 36, (3) although the median ratio of 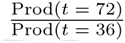 is close to 1 (indicating perfect stability for the median sample, something difficult to achieve), the mean ratio is significantly higher than 1, indicating some samples have very high productivities at *t* = 72, and (4) there are some 2-fold more highly productive samples at *t* = 72 than at *t* = 36. These and other patterns indicate there are issues with the numerical relative productivity data from Ref. [39]. To mitigate these problems, the numerical data were converted to classes (as described in Section 2.1).

**Table A1:** Analysis of the numerical relative productivity data from Ref. [39]. As mentioned in Section 2.1, a sample was labeled as “stable” if its relative productivities after 36 and 72 population doublings were *≥* 0.7; else, it was labeled as “unstable”. For the 3-class problem, a sample was labeled as “highly productive” if its relative productivities after 36 and 72 population doublings were *≥* 2. Percentages are relative to the number of samples for each dataset.

| Propriety | Train Set | Test Set |
| --- | --- | --- |
| Number of samples | 6,301 (100%) | 1,341 (100%) |
| Stable (both periods) | 4,113 (65.3%) | 920 (68.6%) |
| Stable ($t = 36$) | 5,268 (83.6%) | 1,117 (83.3%) |
| Stable ($t = 72$) | 4,477 (71.1%) | 1,021 (76.1%) |
| $\text{Prod}(t = 36) > \text{Prod}(t = 72)$ | 3,052 (48.4%) | 695 (51.8%) |
| Mean and Median $\frac{\text{Prod}(t = 72)}{\text{Prod}(t = 36)}$ ratio | 1.51; 1.022 | 1.38; 0.979 |
| Highly Productive (both periods) | 323 (5.13%) | 45 (3.36%) |
| Highly Productive ($t = 36$) | 493 (7.82%) | 89 (6.64%) |
| Highly Productive ($t = 72$) | 1,087 (17.3%) | 164 (12.2%) |

## Appendix A3 Methylation data do not improve the predictive performance of models

As per Section 2.1, the raw epigenetic data of Ref. [19] also contained methylation data, which we processed for model training. The number of methylated residues in and surrounding a potential hotspot was included in a one-hot encoded format. Multiple window sizes are tested, such that a window size of X includes the 16 hotspot residues plus X residues before and after the hotspot. As mentioned in Section 3.1 and shown in Fig. 2, the inclusion of these methylation data does not improve the predictive performance of the models. Moreover, the use of only methylation data leads to poorly performing models. We hypothesize that this lack of effect may also be due to the small percentage of methylated residues, even when a large window size is used (Table A2). Models trained with chromatin and methylation data (Table A3 and Fig. A1) have similar, and statistically indistinguishable, F1 scores and MCCs to models trained with only chromatin data (Table 1 and Fig. 1). Models trained with only methylation data have low performance (Table A4). As such, only the chromatin data were used to train the models in the main text.

**Table A2:**
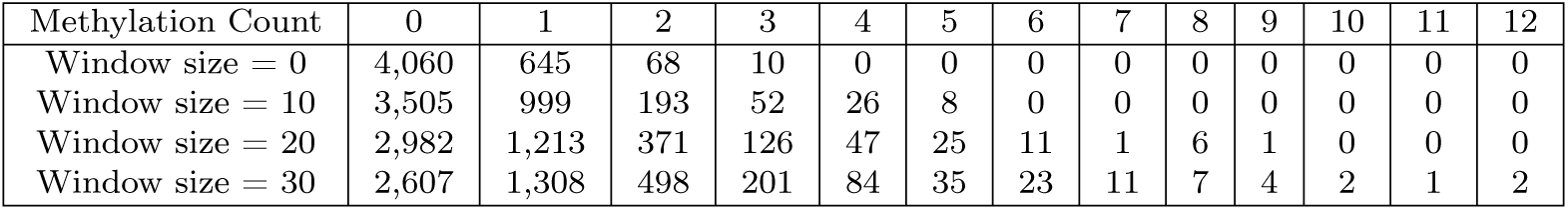
Number of training-set samples with a certain methylation count for different window sizes. For reference, the preprocessed training set contains 4,783 samples.

| Methylation Count | 0 | 1 | 2 | 3 | 4 | 5 | 6 | 7 | 8 | 9 | 10 | 11 | 12 |
| --- | --- | --- | --- | --- | --- | --- | --- | --- | --- | --- | --- | --- | --- |
| Window size = 0 | 4,060 | 645 | 68 | 10 | 0 | 0 | 0 | 0 | 0 | 0 | 0 | 0 | 0 |
| Window size = 10 | 3,505 | 999 | 193 | 52 | 26 | 8 | 0 | 0 | 0 | 0 | 0 | 0 | 0 |
| Window size = 20 | 2,982 | 1,213 | 371 | 126 | 47 | 25 | 11 | 1 | 6 | 1 | 0 | 0 | 0 |
| Window size = 30 | 2,607 | 1,308 | 498 | 201 | 84 | 35 | 23 | 11 | 7 | 4 | 2 | 1 | 2 |

**Table A3:**
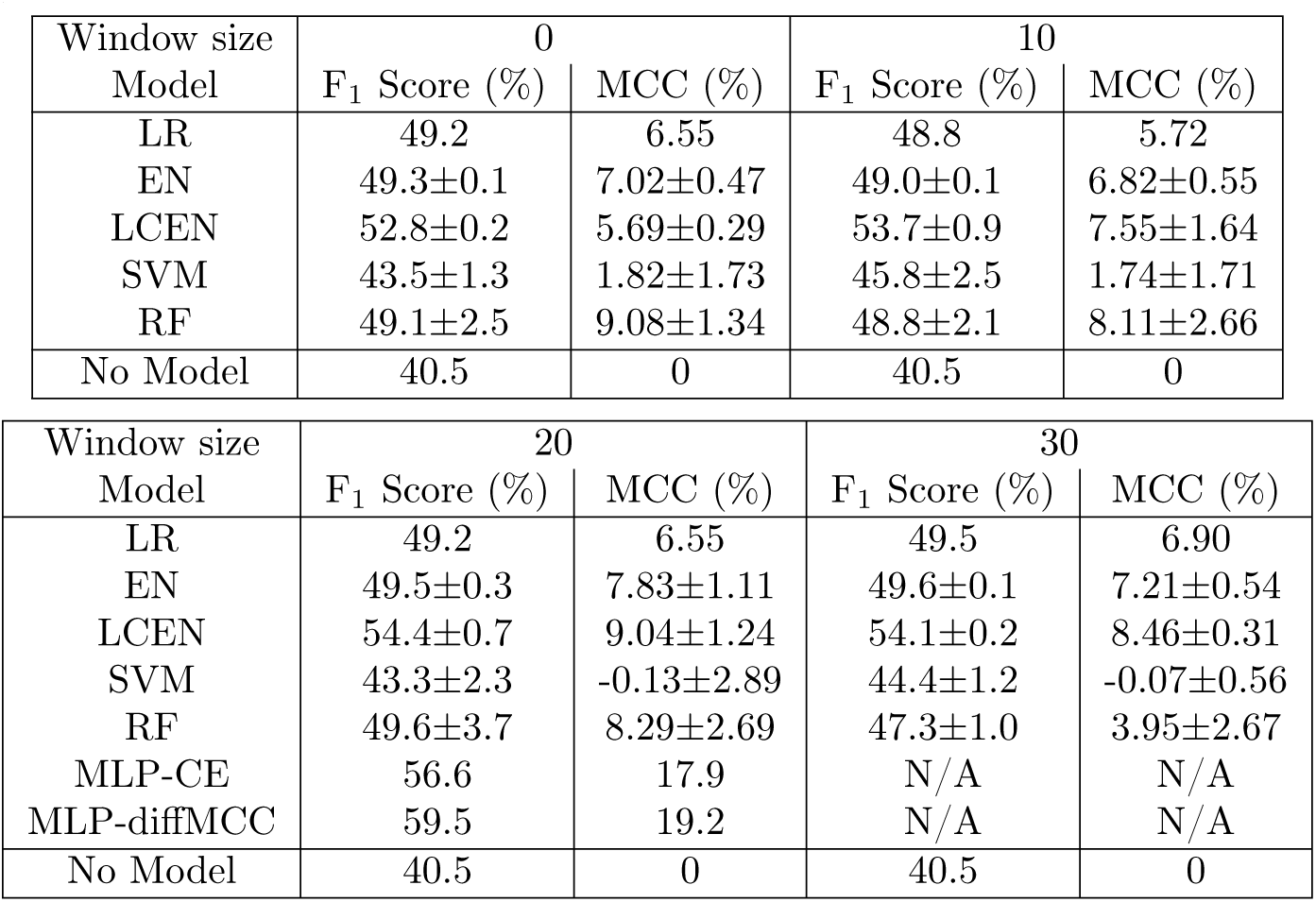
Test-set averaged F_1_ scores and MCC for different models trained with the 2-class version of the stability data as a function of chromatin modification and methylation levels. Model labels are as in Section 2.2. “No Model” refers to the F_1_ score obtained when all samples are guessed as positive and is the minimum value any model should achieve. Compare with Table 1. Some MLP models were not trained in the interest of time due to the models’ unchanged performance (labeled as “N/A” here).

## Appendix A4 Computational resources used

All experiments were done in a personal computer equipped with a 13th Gen Intel^®^ Core™ i5-13600K CPU, 64 GB of DDR4 RAM, and an NVIDIA GeForce RTX 4090 GPU.

---

**Fig. A1:**
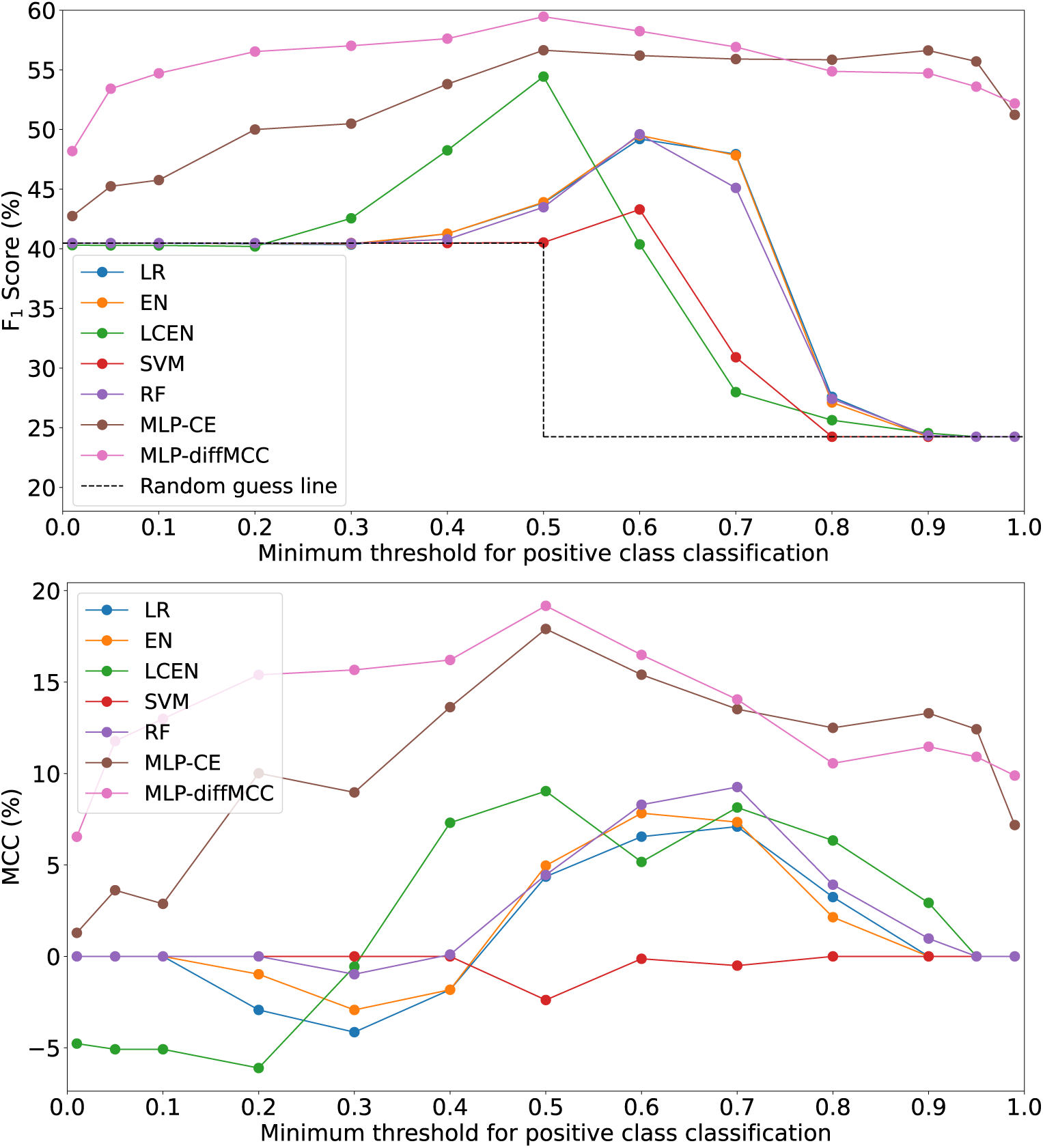
Averaged F_1_ scores (top) and Matthew correlation coefficients (MCCs) (bottom) for the models trained with the 2-class version of the stability data as a function of chromatin modification and methylation levels with a window size of 20. Model labels are as in Section 2.2. “Random guess line” shows the F_1_ score obtained by guessing all samples as positive or negative. Compare with Fig. 1.

**Table A4:**
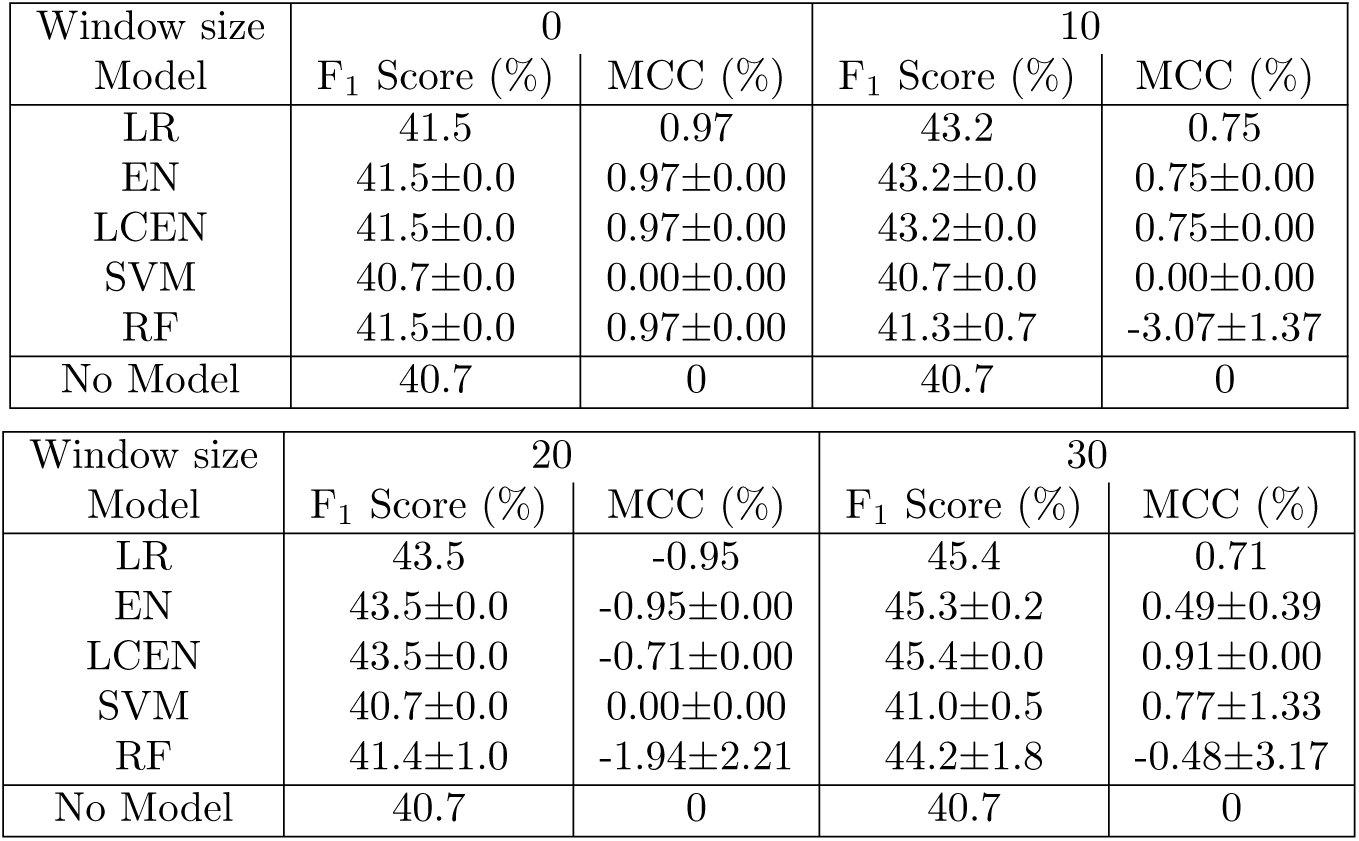
Test-set averaged F_1_ scores and MCC for different models trained with the 2-class version of the stability data as a function of only methylation levels. Model labels are as in Section 2.2. “No Model” refers to the F_1_ score obtained when all samples are guessed as positive and is the minimum value any model should achieve. Compare with Table 1. MLP models were not trained in the interest of time due to the models’ low performance.

| Window size<br>Model | 0 |  | 10 |  |
| --- | --- | --- | --- | --- |
| | $F_1$ Score (%) | MCC (%) | $F_1$ Score (%) | MCC (%) |
| LR | 41.5 | 0.97 | 43.2 | 0.75 |
| EN | $41.5 \pm 0.0$ | $0.97 \pm 0.00$ | $43.2 \pm 0.0$ | $0.75 \pm 0.00$ |
| LCEN | $41.5 \pm 0.0$ | $0.97 \pm 0.00$ | $43.2 \pm 0.0$ | $0.75 \pm 0.00$ |
| SVM | $40.7 \pm 0.0$ | $0.00 \pm 0.00$ | $40.7 \pm 0.0$ | $0.00 \pm 0.00$ |
| RF | $41.5 \pm 0.0$ | $0.97 \pm 0.00$ | $41.3 \pm 0.7$ | $-3.07 \pm 1.37$ |
| No Model | 40.7 | 0 | 40.7 | 0 |

| Window size<br>Model | 20 |  | 30 |  |
| --- | --- | --- | --- | --- |
| | $F_1$ Score (%) | MCC (%) | $F_1$ Score (%) | MCC (%) |
| LR | 43.5 | -0.95 | 45.4 | 0.71 |
| EN | $43.5 \pm 0.0$ | $-0.95 \pm 0.00$ | $45.3 \pm 0.2$ | $0.49 \pm 0.39$ |
| LCEN | $43.5 \pm 0.0$ | $-0.71 \pm 0.00$ | $45.4 \pm 0.0$ | $0.91 \pm 0.00$ |
| SVM | $40.7 \pm 0.0$ | $0.00 \pm 0.00$ | $41.0 \pm 0.5$ | $0.77 \pm 1.33$ |
| RF | $41.4 \pm 1.0$ | $-1.94 \pm 2.21$ | $44.2 \pm 1.8$ | $-0.48 \pm 3.17$ |
| No Model | 40.7 | 0 | 40.7 | 0 |

1 e.g.: *β* = 3 indicates recall is 3 times as important as precision; *β* = 0.5 indicates precision is 2 times as important as recall.

2 Equivalent to selecting the 5% most productive samples.

## Notes

https://github.com/PedroSeber/CHO_stability_prediction

### Competing Interest Statement

The authors have declared no competing interest.

### Summary of Updates

Added Fig. 5, clarified methodology/data preprocessing, and expanded on the Discussion section.

